# Computational framework for targeted high-coverage sequencing based NIPT

**DOI:** 10.1101/486282

**Authors:** Hindrek Teder, Priit Paluoja, Andres Salumets, Kaarel Krjutškov, Priit Palta

## Abstract

Non-invasive prenatal testing (NIPT) enables accurate detection of fetal chromosomal trisomies. The majority of existing computational methods for sequencing-based NIPT analyses rely on low-coverage whole-genome sequencing (WGS) data and are not applicable for targeted high-coverage sequencing data from cell-free DNA samples.

Here, we present a novel computational framework for a targeted high-coverage sequencing based NIPT analysis. The developed methods use a hidden Markov model (HMM)-based approach in conjunction with supplemental machine learning methods, such as decision tree (DT) and support vector machine (SVM), to detect fetal trisomy and parental origin of additional fetal chromosomes. These methods were tested with simulated datasets covering a wide range of biologically relevant scenarios with various chromosomal quantities, parental origins of extra chromosomes, fetal DNA fractions and sequencing read depths. Consequently, we determined the functional feasibility and limitations of each proposed approach and demonstrated that read count-based HMM achieved the best overall classification accuracy of 0.89 for detecting fetal euploidies and trisomies. Furthermore, we show that by using the DT and SVM methods on the HMM state classification results, it was possible to increase the final trisomy classification accuracy to 0.98 and 0.99, respectively.

We demonstrated that read count and allelic ratio-based models can achieve a high accuracy (up to 0.98) for detecting fetal trisomy even if the fetal fraction is as low as 2%. Currently existing methods require at least 4% fetal fraction, which can be an issue in the case of early gestational age (<10 weeks) or elevated maternal body mass index (>35 kg/m^2^). More accurate detection can be achieved at higher sequencing depth using HMM in conjunction with supplemental methods, which significantly improve the trisomy detection especially in borderline scenarios (e.g., very low fetal fraction) and can enable to perform NIPT even earlier than 10 weeks of pregnancy.

## Introduction

It is well known that chromosomal aneuploidies are the leading cause of spontaneous miscarriages and congenital disorders in humans (1,2). At least 10% of all clinically diagnosed pregnancies are trisomic or monosomic. It is assumed that many aneuploid conceptions are eliminated during the earliest stages of pregnancy (3). The most common aneuploidies are trisomies, which are characterized by the presence of an additional chromosome and caused by segregation errors, occurring during meiotic divisions. In case of trisomy of chromosome 21, approximately 90% are of maternal origin and 73% occur during first meiotic division (4–9). Despite routinely performed prenatal screenings in most developed countries, more than 0.1% of all live births are trisomic and the corresponding risk continues to rise with increasing maternal age (10).

Advanced non-invasive methods for prenatal screening using cell-free DNA (cfDNA) have considerably improved the detection of fetal aneuploidies (11). The most commonly used technique, whole-genome sequencing (WGS)-based non-invasive prenatal testing (NIPT) enables inference of the ploidy of each chromosome by counting the specifically mapped sequencing reads to each chromosome (12,13). Although NIPT offers increased accuracy compared to the first trimester serum screening and ultrasound, it is usually not a part of conventional prenatal screenings due to its high cost.

Alternative NIPT techniques have the potential to reduce high-cost limitations by using a targeted sequencing approach (14–16). Instead of low coverage WGS, only certain genomic regions are analyzed at high coverage. Targeting involves the use of hybridization-based capture or multiplex PCR amplification to enrich the genomic regions of interest (14,15). Compared to the WGS-based methods, targeted approaches require less cfDNA and enable to study more samples in parallel, making it a cost-efficient alternative. A few already available targeted solutions rely on sequencing single nucleotide polymorphisms (SNPs). In these cases, allelic information from sequencing read counts can be used to calculate allelic ratios obtained from heterozygous SNPs and also serve as an extra source of information for inferring fetal aneuploidies (17). For example, NATUS software, developed by Natera, Inc., considers parental genotypes and crossover frequency data to calculate the expected allele distributions for SNPs and possible fetal genotypes based on recombination sites in the parental chromosomes (18). The algorithm compares predicted allelic distributions with measured allelic distributions by employing a Bayesian-based maximum likelihood approach to determine the relative likelihood of chromosomal copy number hypothesis. The likelihoods of each sub-hypothesis are summarized and the hypothesis with the maximum likelihood is the chromosome copy number in the fetal DNA fraction (FF). Although feasible, this method is proprietary and not available to the community. An alternative approach is to model a chromosome as hidden Markov model (HMM) of sequential loci and determine the most likely chromosomal copy number status at each locus and consequently the overall chromosomal ploidy. Kermany and colleagues used HMM to detect fetal trisomy using high-density SNP markers from a trisomic individual and one parent (19), and similar HMM-based approaches have been previously used to detect both full and sub-chromosomal aneuploidies using binned read counts (20,21).

In the current study, we present a novel statistical framework for detecting fetal trisomy and possibly the parental origin of the trisomy from targeted high-coverage sequencing data of pregnant women’s cfDNA. The framework incorporates three different HMMs that utilize read counts of targeted loci, allelic ratios of targeted SNPs, or both in combination with a decision tree (DT) or support vector machine (SVM)-based trisomy detection, without requiring any prior knowledge of parental genotypes. We provide a comprehensive evaluation of the performance and limitations of these methods on simulated datasets generated for a wide range of biologically and technically relevant scenarios. These results can be used as guidelines for appropriate study design and feasibility analysis for future NIPT studies using targeted sequencing approach.

## Materials and Methods

### Sequencing data simulation

A total of 1,800 datasets were generated with different parameters to mimic the read count data obtained from targeted sequencing of 10,000 pregnant women’s cfDNA samples in various conditions. Simulated datasets varied in the context of (1) fetal condition – euploidy, maternally or paternally originated trisomy characteristic to meiosis I segregation failure; (2) sequencing read depth (RD) – in the range of 500 to 15,000 at increments of 500; and (3) FF – in the range of 1 to 20% at increments of 1%. Each dataset incorporated 10,000 individual chromosome sets, each chromosome incorporated 1,000 SNPs.

As the cfDNA of a pregnant woman contains both maternal and fetal DNA, we started the simulation with the formation of parental chromosomes. For both parents, we generated two sets of 1,000 SNPs representing a pair of homologous chromosomes. Each SNP was biallelic and both alleles had an equal likelihood of occurrence (MAF = 0.5). Before creating a fetal set of chromosomes, parental homologous chromosomes underwent a chromosomal crossover by exchanging a random number of homologous alleles. The resulting recombined chromosomes were used to form a set of fetal chromosomes according to the fetal conditions.

In addition, we generated allele counts for each SNP according to the mean sequencing coverage and FF of the dataset. One might assume that all reads in a given region would follow a Poisson distribution with a mean proportional to the copy number of the region. However, due to the various technical biases, the process is over-dispersed and the simulation distribution followed the negative binomial distribution with a variance-to-mean ratio of 3 (22).

### Allelic ratio calculation

Based on the simulated data, we calculated the allelic ratio for every “informative” SNP. Only SNPs which were heterozygous in mother and/or fetus were considered as informative. If both alleles have equal likelihood of occurrence (MAF = 0.5), on average 75% of SNPs were informative in case of maternally originated trisomy and the proportion of informative SNPs was even higher in the case of paternally originated trisomy as both paternal alleles contributed to heterozygosity independently. The allelic ratio was defined as the number of sequencing reads carrying a major allele for a certain variant divided by the number of sequencing reads carrying a minor allele.

### Fetal fraction calculation

FF showed the proportion of fetal cfDNA in total cfDNA. We estimated the FF of a cfDNA sample using the allelic counts of the sample’s reference chromosome. First, we filtered the informative SNPs on the reference chromosome, where the mother was homozygous and the fetus was heterozygous (allelic ratio > 2.5). In this subset, the major allele count was the sum of maternal allele counts and 1/2 of the fetal allele count. The minor allele count was proportional to 1/2 of the fetal allele count. The FF was calculated as the median value of the ratios between 2 × minor allele counts and the sum of major and minor allele counts.

The FF of a sample was calculated using the following formula:

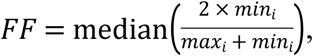

where FF denotes the fetal fraction, max_i_ – the major allele count of SNP *i*, and min_i_ – the minor allele count of SNP *i.* The median value over all informative SNPs was considered as estimated FF of a sample, which showed high similarity to actual FF (Fig in S2 Fig).

### Hidden Markov model

For the detection of fetal trisomy and the parental origin of the trisomy, we implemented HMM in Python (version 3.6.2) using the hmmlearn (version 0.2.0) package. First, we created three distinct models based on the observed measurements of sequential SNPs – (1) read counts (Fig A in S1 Fig), (2) allelic ratios (Fig B in S1 Fig), and (3) the combination of both read counts and allelic ratios (Fig B in S1 Fig). Second, we estimated the parameters for the models empirically using a simulated training dataset. Finally, we used the Viterbi algorithm to find the most likely underlying fetal condition behind each SNP.

### Read count model

The read count (RC) model is a 2-state HMM which enables detection of underlying fetal conditions of sequential SNPs using read counts (Fig A in S1 Fig). The possible outcome states of the model are “euploidy” and “trisomy”. The RC model is based on the hypothesis that the mean coverage of a given region is proportional to the copy number of the region. In the case of fetal trisomy, there is an extra chromosome and therefore we would expect to see a 1/3 increase in fetal read counts compared to the euploid chromosome.

### Allelic ratio and combined models

The allelic ratio (AR) model and the combined model of read count and allelic ratio (RCAR) are both 7-state HMMs, which enable detection of underlying fetal conditions and the parental origin of SNPs (Fig B in S1 Fig). The AR model uses allelic ratios of sequential informative SNPs as inputs. The RCAR model incorporates sequential read counts and allelic ratios as inputs. Both models classify loci into seven categories by the allelic pattern. The allelic pattern depends on the maternal and fetal genotypes and the fetal condition (Table in S6 Table). The possible outcome states of the model are “euploidy”, “trisomy”, and “paternal trisomy”. Although the “trisomy” condition includes loci typical to both maternally and paternally originated trisomy, here we associated “trisomy” with maternally originated trisomy to avoid over-estimation of paternally originated trisomy.

### Parameter estimation

In all three HMMs, no prior distribution of the initial state was assumed. Each possible state had an equal likelihood of occurrence. The HMM transition probability was set to 10 times more likely to stay in the same state than to switch between states with different fetal conditions. The emission probabilities were obtained using the training datasets. For each test dataset, we simulated a training dataset of 100 cfDNA samples with corresponding FF and sequencing coverage. In our models, the emission probabilities were approximated to a Gaussian distribution. The distribution parameters were obtained for each state by calculating the mean and variance of the read counts and allelic ratios of the training dataset.

### Fetal condition estimation

The chromosomal condition of a cfDNA sample was determined by the most frequently occurring underlying condition of targeted loci using the RC, AR, and RCAR models. If no condition was prevalent, the cfDNA sample was marked as unclassified.

To improve the accuracy, especially in the case of paternally originated trisomy, we applied the DT and the SVM on HMM-classified state proportions of the targeted loci. Both methods were implemented in Python (version 3.5.5) using scikit-learn (version 0.19.1). The DT was used with default parameters, except the maximum depth of the tree was set to three and the random state generator to 123. The SVM also used default parameters and the random state generator was set to 123. As the DT and SVM are supervised learning models, we used the training dataset to fit the models. Eventually, each cfDNA sample was classified using both models by the following features – RD, FF and HMM state frequencies. The possible classification output values were identical to HMM.

## Results and Discussion

We developed three novel HMM-based statistical methods to detect fetal chromosomal trisomies from targeted sequencing assays. In addition to a naïve HMM-based frequentist approach for trisomy detection, we applied two machine learning (ML) methods to infer fetal trisomy. While considering a wide range of biologically and technically motivated conditions, we simulated datasets mimicking cfDNA sequencing assays and used these data to perform a comprehensive evaluation of our proposed computational methods (Fig 1).

### Novel HMM-based methods for trisomy detection

By considering the sequencing read counts (RC) of targeted loci, allelic ratios (AR) of targeted SNPs, or both (RCAR), the developed HMM models were used to classify consecutive target loci on a studied chromosome into pre-defined underlying states. In the 2-state RC model, these unique states represented fetal euploidy and trisomy (Fig A in S1 Fig). In the case of the 7-state AR and RCAR models, these different states can occur with fetal euploidy or maternally/paternally originated trisomy (Fig B in S1 Fig). Consequently, the proportion of loci classified into these distinct states can be used to estimate the fetal condition of each studied chromosome (see “Fetal condition estimation” in Methods). And although such naïve classification works relatively well in case of high sequencing read depth (RD) and fetal fraction (FF) scenarios, the proportion of loci classified into these underlying states can be similar and thus difficult to distinguish unambiguously in the case of low RD and FF (Fig 2).

Therefore, the precise calculation of FF is also crucial for controlling the precision and uncertainty of fetal trisomy detection and sequencing-based NIPT. Notably, in the case of the RC model and autosomal chromosomes there is no information that could be used to infer the FF of the studied sample so that optimal corresponding model parameters can be used. One possible solution to overcome this challenge is to use the expected median FF of 10% (23). In the case of the AR and RCAR models, we used informative polymorphic SNPs with heterozygous alleles in mother and/or fetus to infer the sample-specific FF (Fig in S2 Fig), similarly to previous studies (24–26). Additionally, in the case of the AR and RCAR models, allelic count data at informative SNPs can be used to calculate allelic ratios, distinguishing maternally and paternally originated trisomies (see “Allelic ratio calculation” in Methods) according to their distinct allelic patterns (Table in S6 Table). On the other hand, these models only consider informative targeted SNPs that are polymorphic in a given sample, which reduces the total number of analyzed SNPs least by 25% and therefore somewhat decreases the detection accuracy (data not shown).

### Supplemental methods for trisomy detection

Since in some possible scenarios, such as paternally originated trisomy, the previously described HMM-based models did not unambiguously infer the underlying fetal condition (Fig 2), we developed two additional “supplemental” machine learning (ML)-based methods to improve the sample classification accuracy. The supplemental methods, which take HMM-classified state proportions as input, significantly improved the sample classification especially when the proportion of loci inferred into one or the other HMM state was not an obvious majority and where the frequentist approach, therefore, did not work (Table 1 and 2).

All three HMMs (RC, AR, and RCAR) independently and conjointly with the supplemental methods (DT and SVM) were tested on the same collection of simulated cfDNA datasets representing all combinations of different fetal chromosomal conditions (euploidy, maternally and paternally originated trisomy) and FFs (1-20%) sequenced with various RDs (500-15,000 reads), which is feasible for targeted sequencing assays.

### Read count (RC) model

The RC model enables detection of fetal euploidy and trisomy by using sequencing read counts in successive (targeted) regions along the chromosome of interest. As read count data alone cannot be used to infer the FF of a studied sample, we assumed FF as 10% in this testing model. Nevertheless, the HMM method showed excellent accuracy in detecting fetal euploidy (Fig 3). On the other hand, this method was ineffective at detecting fetal trisomy if the FF was lower than 6% and increasing the RD induced only a minor increase in detection accuracy (Table 1). It is also important to note that since there is no direct method to distinguish between paternally and maternally inherited alleles, the read count model does not enable determination of the parental origin of the trisomy. Since it uses only sequencing read count information to detect fetal trisomies, it is relatively straightforward to integrate this model with most existing sequencing-based solutions.

In general, applying supplemental methods significantly improved the RC model-based classification at lower FFs (Table 2). The DT method allowed accurate detection of fetal euploidy and trisomy even if the FF was as low as 3%; the SVM method successfully lowered that limit even further, allowing accurate detection of fetal trisomies at FF 2%, with a small trade-off in detecting aneuploid chromosomes (Fig 3). Unexpectedly, DT trisomy detection improved at a lower read coverage. This can be explained by the strictly set maximum depth (*max_depth* = 3) of the DT, which prevented overfitting of the model; on the other hand, this method was not suitable for classifying a wide range of FF values. This shortcoming is due to the fixed FF parameter rather than the properties of the DT (Fig 3, Fig in S3 Fig).

### Allelic ratio (AR) model

The AR model uses counts of sequencing reads containing one or the other allele at informative SNP loci along the chromosome of interest to estimate if the studied sample has euploid, maternally or paternally originated trisomy to infer the FF of the corresponding sample. The AR model showed excellent accuracy detecting fetal euploidy even at an FF of 1% and an RD of 500 and reasonable accuracy to detect maternally originated trisomy if FF was ≥ 6% and RD was higher than 10,000 (Fig 4). In contrast to the DT and the SVM methods, it was unable to detect paternally originated trisomy in a given range of FF and RD (Fig in S4 Fig).

Compared to the read count data, allelic ratio information was used to estimate the FF of a sample using specific allelic patterns (Table in S6 Table). In addition, allelic ratio data were used to separate maternally and paternally originated trisomies. As for the HMM, the inability to detect paternally originated trisomy can be explained by the overlapping emission distributions of the allelic ratios of maternally and paternally originated trisomies.

In general, the supplementary methods increased the detection accuracy for the AR model significantly (Table 2), especially in the case of paternally originated trisomy (Table 1, Fig in S4 Fig). In the case of maternally originated trisomy, all three methods had similar characteristics as the detection accuracy was positively correlated with both sequencing RD and FF (Fig 4). The read count had a stronger impact on the AR model, whereas the RC model was mostly affected by FF. The DT had a slight fetal trisomy detection improvement compared to the HMM, and the SVM in turn had a slight advantage over the DT. DT methods also showed excellent accuracy in detecting fetal euploidy. Unlike the other methods, the SVM showed slightly better maternally originated trisomy detection accuracy and consistently good results if the read coverage was low (RD = 500); on the other hand, the SVM had poor results detecting fetal euploidy if the read coverage was low (RD = 500). The SVM failure for euploidy and excellent results for maternally originated trisomy at low read coverage contradicted each other, which was a sign of maternally originated trisomy over-estimation. In the case of paternally originated trisomy, the DT and SVM had excellent detection accuracy (Table 1).

### Combined (RCAR) model

Finally, we studied the RCAR model, which incorporates both read count and allelic ratio information to predict fetal euploidy or trisomy. Furthermore, it utilizes informative SNPs, which enables separation of maternally and paternally originated trisomy by allelic patterns (Table in S6 Table) and estimated FF. The RCAR model showed excellent results in detecting fetal euploidy (Fig in S5 Fig). Compared to the HMM, the supplemental methods were inefficient to detect fetal euploidy when the FF and read coverage were low (RD ≤ 1,500; FF ≤ 3%). All three methods showed a positive correlation between detection accuracy, RD and FF, while the HMM detection accuracy was approximately twice as worse compared to the supplemental methods. In case of maternally originated trisomy, the DT and the SVM had better detection accuracy than HMM (Fig 4). In the case of paternally originated trisomy, the DT had excellent detection accuracy followed closely by the SVM (Table 1). However, the HMM was unable to detect paternally originated trisomy in any give range of FF and read coverage (Fig in S5 Fig).

The RCAR model showed significantly higher accuracy in conjunction with supplemental methods (Table 2). Compared to the HMM, the supplementary methods increased the detection accuracy in the case of fetal trisomies (Fig in S5 Fig). As for the HMM, the inability to detect paternally originated trisomy can be explained by the overlapping emission distributions (allelic ratios) of maternally and paternally originated trisomy. Similarly to the AR model, the overall accuracy of the RCAR model was affected by both FF and sequencing RD, whereas the RC model was mostly affected by FF (Fig in S5 Fig, Fig 3).

## Conclusions

Targeted sequencing approaches have the potential to reduce the price of NIPT and improve the quality of healthcare. In the current study, we present HMM-based models in conjunction with supplemental methods (DT and SVM), which enabled the detection of fetal trisomy and the parental origin of an extra chromosome using targeted sequencing-based prenatal (NIPT) assays. The developed methods were tested on simulated datasets generated for a wide range of biologically and technically motivated scenarios to determine the functional feasibility and limitations of each approach.

We determined that regardless of the computational method used, the most challenging factor in fetal trisomy detection is low FF. In our study, the RC model in conjunction with ML-based supplemental methods can detect fetal trisomy at 2% FF, which enables earlier testing compared to the current NIPT assays. Although the RC model can be easily incorporated into currently available targeted workflows, the RCAR model is the recommended choice for its high accuracy and ability to determine the parental origin of the trisomy and to accurately estimate the studied sample FF.

## Acknowledgments

We would like to acknowledge Jüri Lember and Mart Kals for their insights and useful comments. This work was supported by the Estonian Ministry of Education and Research (grant no IUT34-16) and by the Enterprise Estonia (grant no EU48695). Computing was performed at the High Performance Computing Centre of the University of Tartu.

## Supporting information captions

**S1 Fig.**
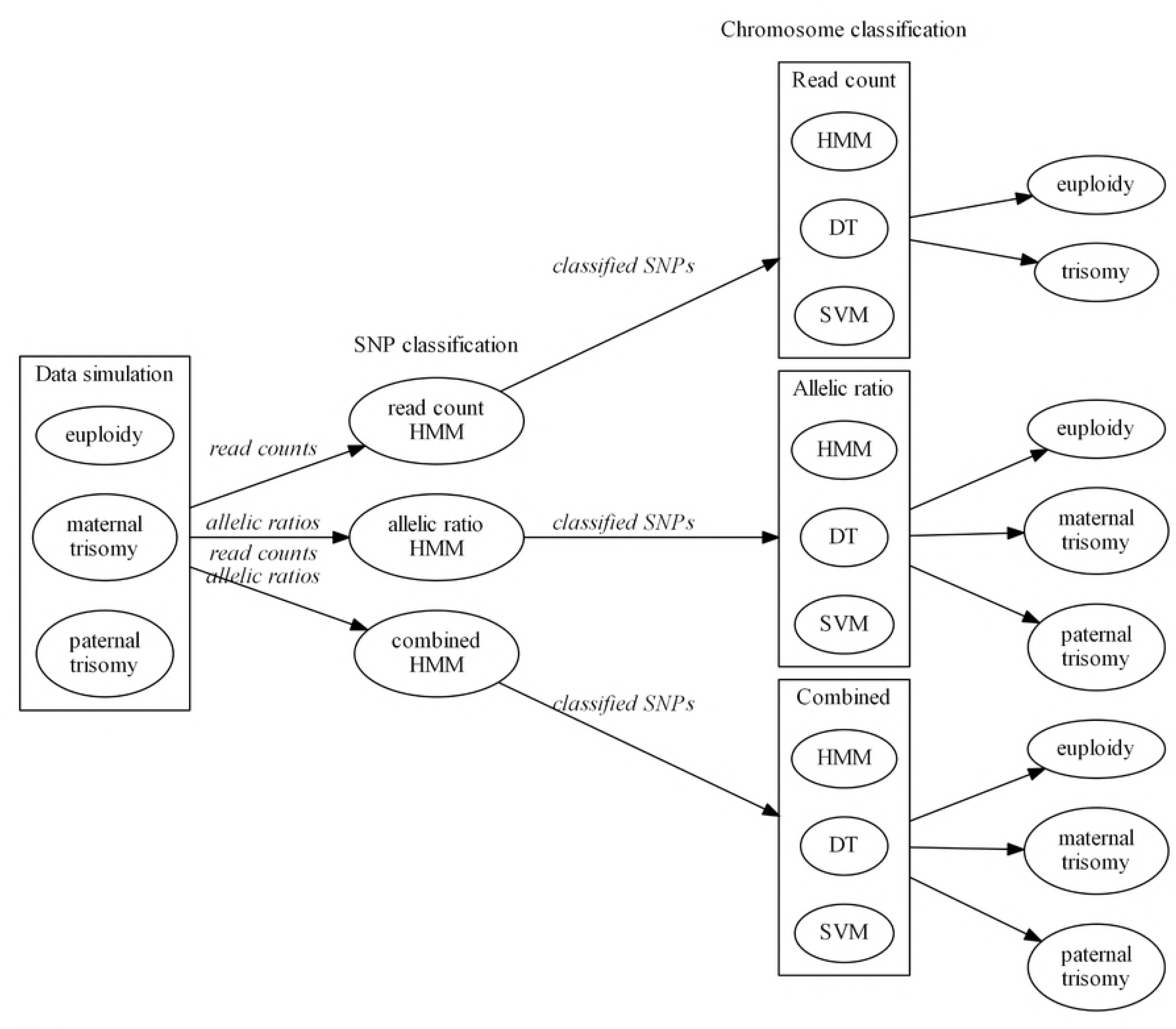
Architecture of 2- and 7-state hidden Markov models (HMMs). (A) The 2-state HMM classified sequential single nucleotide polymorphisms (SNPs) into 2 underlying states, which represent fetal euploidy (white) and trisomy (grey), using read counts. (B) The 7-state HMM classified SNPs into 7 underlying states, which represent fetal euploidy (white), maternally (white-grey) and paternally originated trisomy (grey-white), using allelic ratios with or without read counts.

**S2 Fig.**
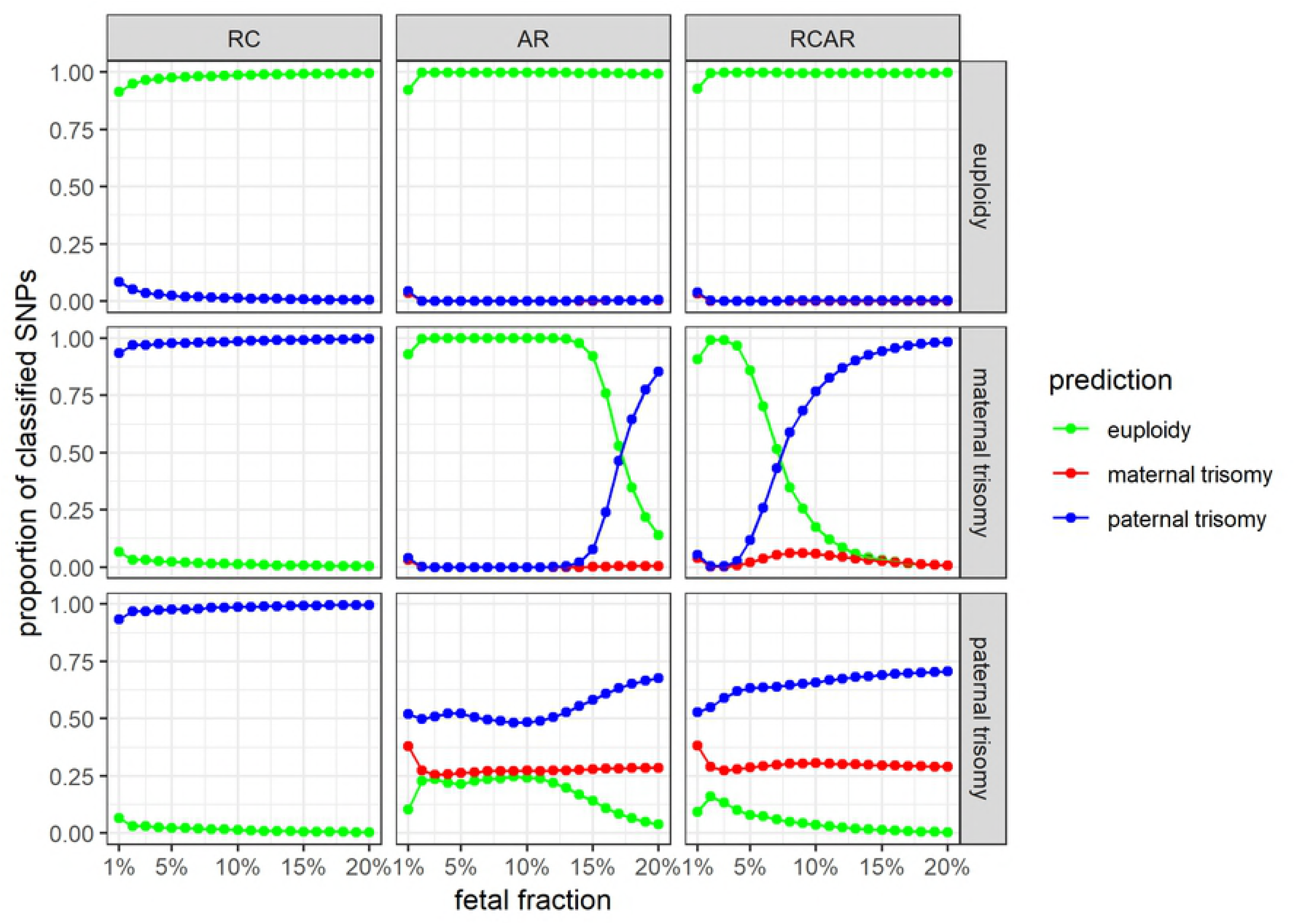
Difference between estimated and simulated fetal fraction (FF). The simulated FF was subtracted from the estimated FF for each simulated cell-free DNA sample to determine the FF difference (y-axis). The differences were grouped as boxplots by sequencing read depth (x-axis). The results show a positive correlation between sequencing read depth and FF estimation accuracy.

**S3 Fig.**
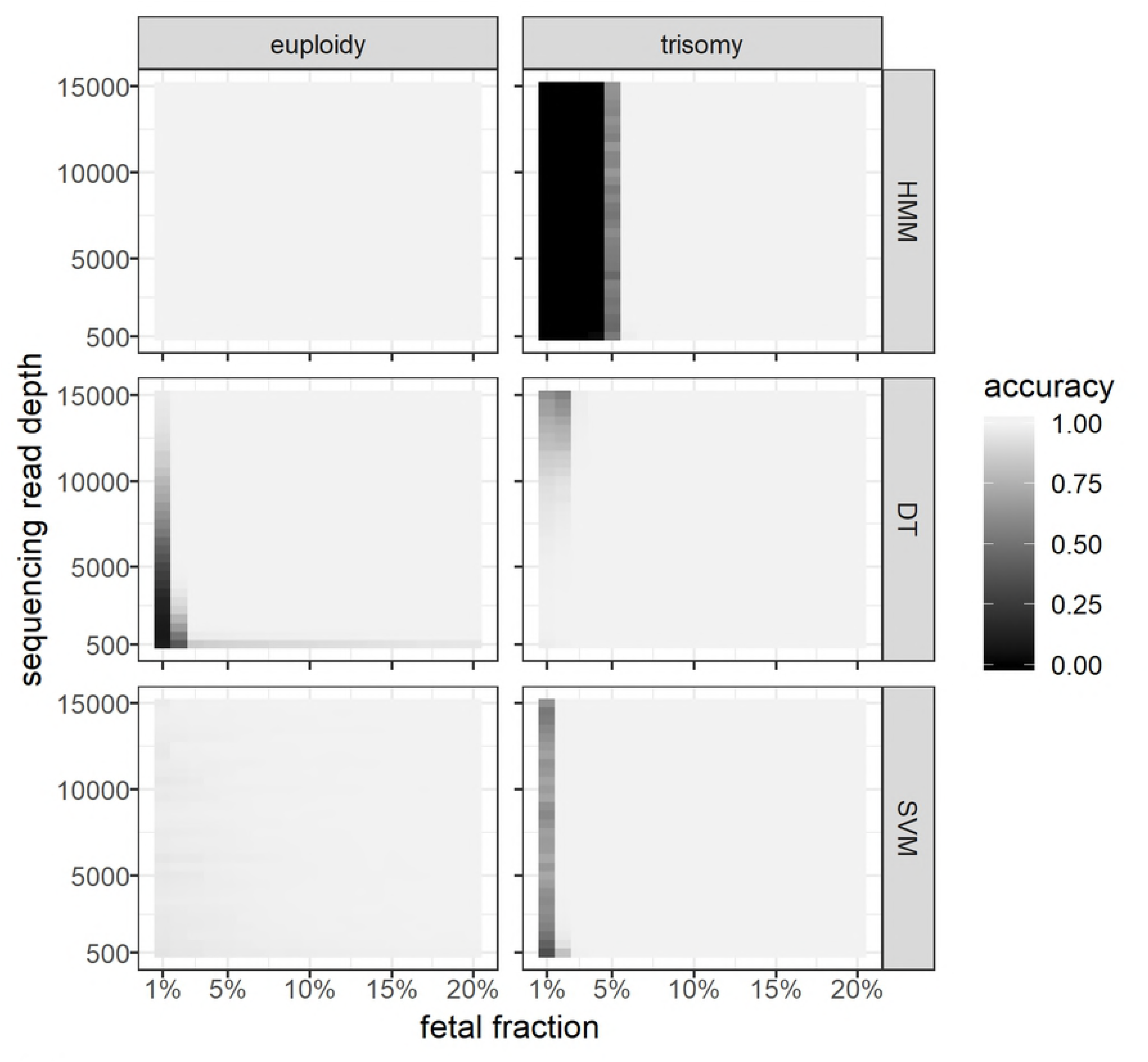
Results of the read count model. The simulated datasets of fetal euploidy and trisomy (vertical panels) were classified by three methods – hidden Markov model (HMM), decision tree (DT) and support vector machine (SVM) (horizontal panels). Each panel includes cells with different fetal DNA fractions (x-axis) and sequencing read coverages (y-axis). Each cell includes 10,000 cell-free DNA samples and the color represents the model classification accuracy.

**S4 Fig.**
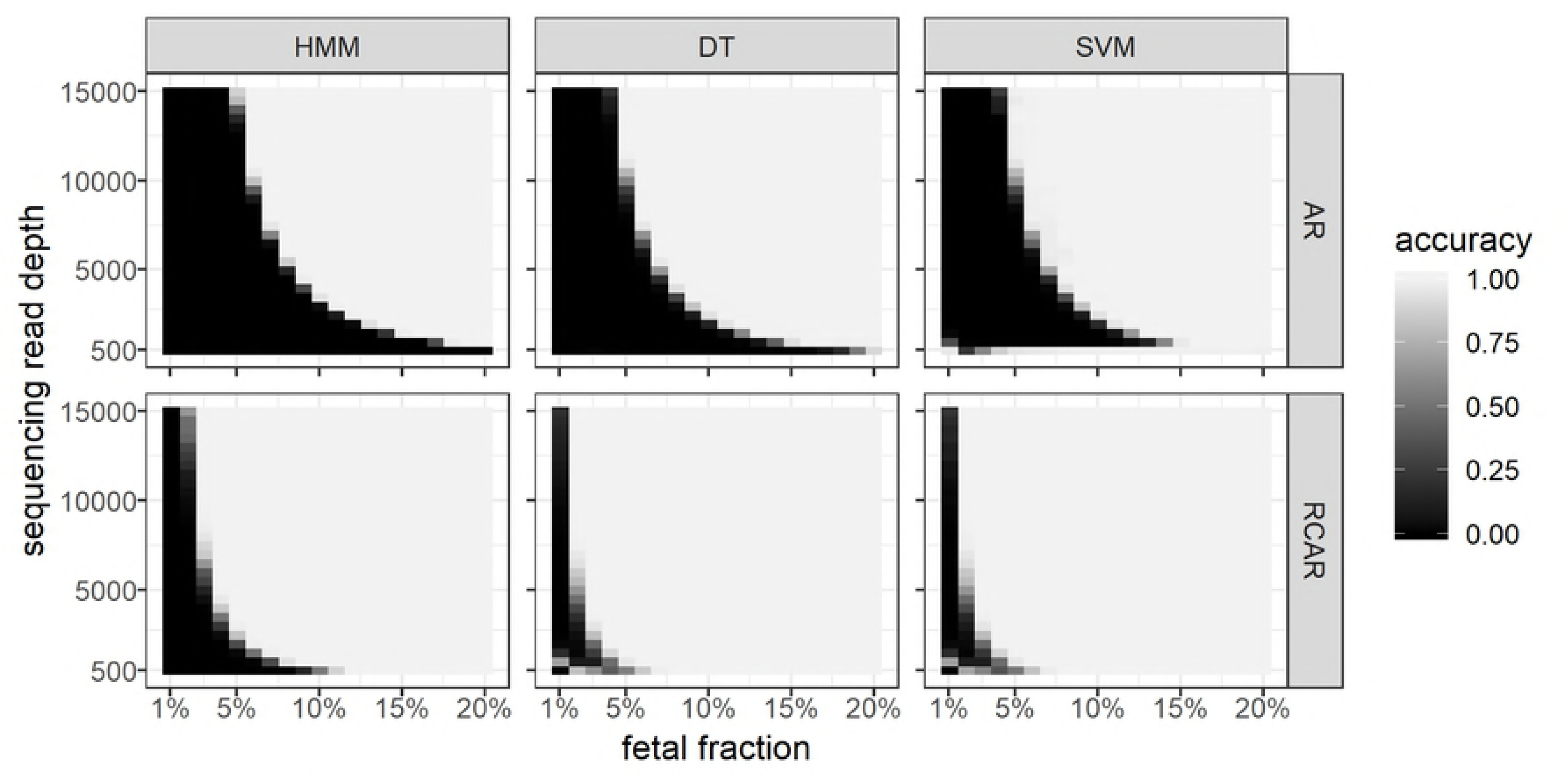
Results of the allelic ratio model. The simulated datasets of fetal euploidy, maternally and paternally trisomy (vertical panels) were classified by three methods – hidden Markov model (HMM), decision tree (DT) and support vector machine (SVM) (horizontal panels). Each panel includes cells with different fetal DNA fractions (x-axis) and sequencing read coverages (y-axis). Each cell includes 10,000 cell-free DNA samples and the color represents the model classification accuracy.

**S5 Fig. Results of the combined model.** The simulated datasets of fetal euploidy, maternally and paternally trisomy (vertical panels) were classified by three methods – hidden Markov model (HMM), decision tree (DT) and support vector machine (SVM) (horizontal panels). Each panel includes cells with different fetal DNA fractions (x-axis) and sequencing read coverages (y-axis). Each cell includes 10,000 cell-free DNA samples and the color represents the model classification accuracy.

**S6 Table. Allelic patterns.** Allelic ratio depends on fetal condition and maternal and fetal genotype.

